# SARS CoV-2 spike adopts distinct conformational ensembles *in situ*

**DOI:** 10.1101/2025.03.04.641425

**Authors:** Amanda J. Gramm, Sean M. Braet, Bindu Y. Srinivasu, Varun Venkatakrishnan, Elijah J. Bass, Fiona L. Kearns, Carla Calvó-Tusell, Rommie E. Amaro, Robert V. Stahelin, Thomas E. Wales, Ganesh S. Anand

## Abstract

Engineered recombinant Spike (S) has been invaluable for determining S structure and dynamics and is the basis for the design of most prevalent vaccines. While these vaccines have been highly efficacious for short-term protection from infection, protection waned with the emergence of variants (alpha through omicron). Here we report differences in conformational dynamics between native, membrane-embedded full-length S and recombinant S. Our virus-like particle (VLP) model mimics the native SARS CoV-2 virion by displaying S assembled with auxiliary E, M, and N proteins in a native membrane environment that captures the entirety of quaternary interactions mediated by S. Display of S on VLP obviates the requirement for stabilizing modifications that have been engineered into recombinant S for enhanced expression and solubility. Amide hydrogen/deuterium exchange mass spectrometry (HDXMS) reveals altered interprotomer contacts in VLP S trimers attributable to the presence of auxiliary proteins, membrane anchoring, and lack of engineered modifications. Our results reveal decreased dynamics in the S2 subunit and at sites spanning interprotomer contacts in VLP S with minimal differences in the N-terminal domain (NTD) and receptor binding domain (RBD). This carries implications for display of epitopes beyond NTD and RBD. In summary, despite affording efficient structural characterization, recombinant S distorts the intrinsic conformational ensemble of native S displayed on the virus surface.

## Introduction

The recent SARS-CoV-2 global pandemic has prompted extensive structural characterization of the SARS-CoV-2 virion. This virion consists of an RNA genome encapsidated by an outer protein shell, which includes the Envelope (E), Membrane (M), Nucleocapsid (N), and Spike (S) proteins (Figure 1A).^1^ The S protein is the largest structural protein that mediates host receptor interactions and facilitates SARS-CoV-2-host cell fusion and entry, making it a primary target for structural biology studies and vaccine design.^2^ The SARS-CoV-2 S is a multifunctional homotrimer, comprised of three protomers that form the characteristic spike structure protruding from the viral lipid bilayer. Each protomer consists of S1 and S2 subunits.^3^ The S1 subunit contains the N-terminal domain (NTD), the receptor-binding domain (RBD), and subdomains 1 and 2 (SD1/2) that contains the RBD hinge.^4^ The S2 subunit encompasses the fusion peptide (FP), two heptad repeats (HR1 and HR2), a transmembrane domain (TM), and cytoplasmic domain (CD) (Figure 1B).^4^ The S1 and S2 subunits are proteolytically processed at position 685, known as the furin cleavage site.^5^ S is responsible for initiating binding to the cellular ACE2 receptor, which requires the RBD to be in the "up" position.^6^ The secondary cleavage event by TMPRSS2 exposes the fusion peptide and initiates cellular membrane fusion and subsequent genomic release.^4,7^

**Figure 1.**
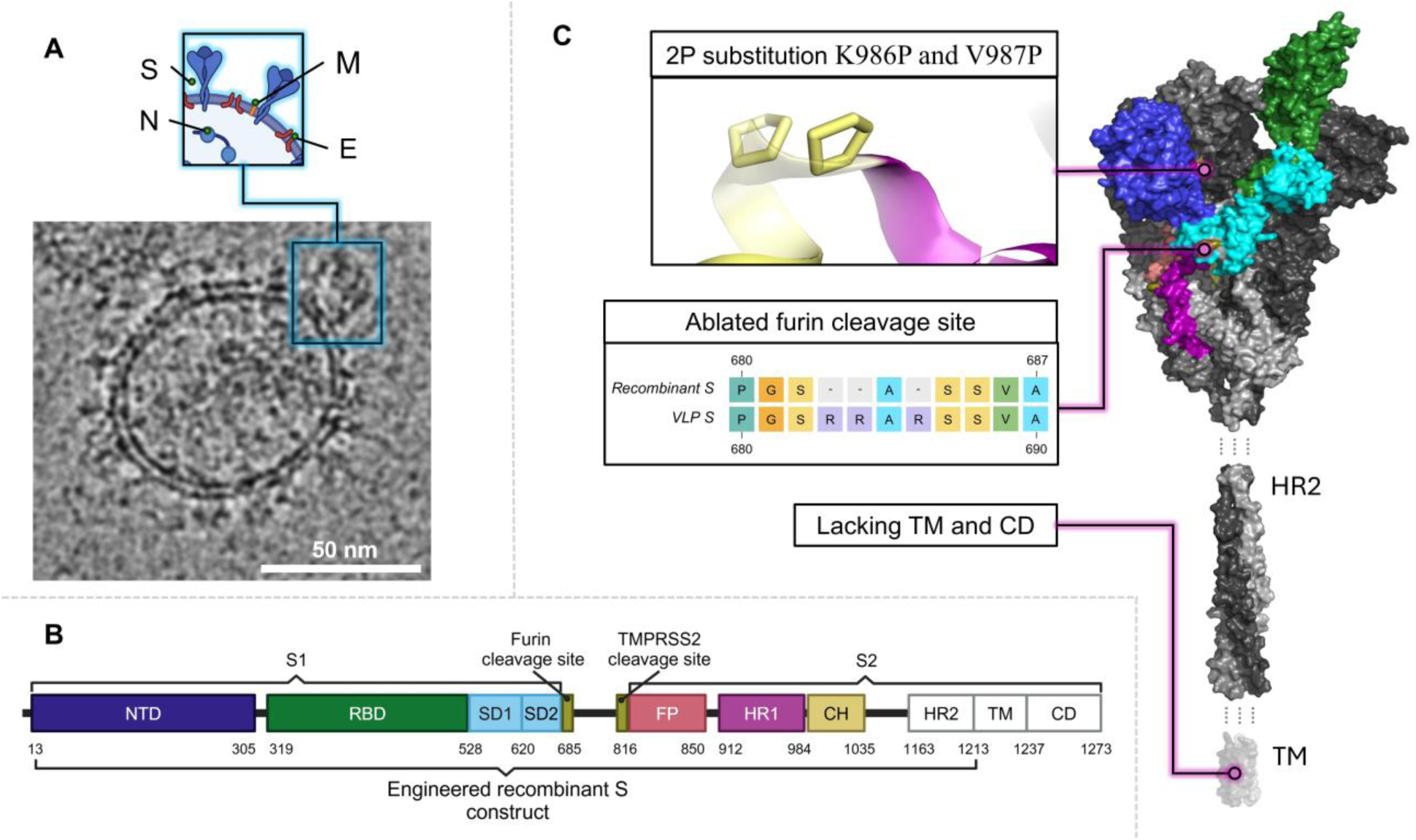
SARS-CoV-2 virion and S trimer domains. **A)** An image of a SARS-CoV-2 VLP with embedded D614G S proteins from a cryo-EM micrograph. An inset illustrating the virion structure provides a cartoon depiction of the organization of structural proteins and lipid bilayer in the native virion (Created using Biorender.com). **B)** Sequence organization of SARS-CoV-2 S. Brackets define the boundaries of the S1 and S2 subunits. The white regions denote domains not included in the recombinant structure (PDB: 7T67). **C)** Structure of D614G S with one RBD in the “up” position (PDB ID: 7T67) indicating engineered differences in recombinant S. The S HR2 domain (PDB: 2FXP) which is unresolved in S structures but present in the recombinant construct and the transmembrane domain (PDB: 7LC8) which is present in the VLP but absent from recombinant S are shown below the spike trimer. The protomers are shown in grayscale, with the NTD, RBD, SD1, SD2, furin cleavage site, TMPRSS2 cleavage site, FP, HR1, and CH are colored according to the sequence organization in B on one protomer.

Constraints posed by sample preparation of the highly infectious SARS-CoV-2 particles have been overcome by generating soluble recombinant S engineered with specific, stabilizing proline substitutions (2P/6P) and an ablated furin cleavage site (Figure 1C).^8,9^ Recombinant S also shows enhanced solubility through removal of the C-terminal membrane anchoring regions and fusion to a T4 trimerization domain. These modifications have enabled high expression of isolated recombinant S trimers to accelerate structural and dynamic analyses.^10,11^ Furthermore, mRNA vaccine constructs, including those from Moderna and Pfizer, have leveraged these strategies to improve post-vaccination expression and stability of S in target human cells for immune recognition.^2^

Previous studies have revealed that S is a highly dynamic trimer that undergoes conformational fluctuations.^12^ This includes RBD up/down dynamics and trimer core splaying in response to cold temperatures.^13^ Additionally, large-scale allosteric changes are observed in the S2 subunit upon ACE2 receptor binding, together with enhanced dynamics near the furin cleavage site.^14^ Mutations in emerging variants have also profoundly impacted dynamics of the recombinant S. Specifically, ACE2 interactions have been attributed to an increased propensity of S in emergent variants to adopt open RBD conformations.^15^ Furthermore, our studies showed that the initial D614G mutation had a significant stabilizing effect on the S trimer, while additional variant mutations increased dynamics across the NTD and RBD.^16^

The recombinant S construct has thus been invaluable for structure determination, epitope mapping, and for mRNA vaccine design. However, the effects of quaternary contacts with auxiliary proteins (E, M, and N) and the membrane bilayer (Figure 1A) on the intrinsic dynamics of native S, have not been fully studied. Additionally, recombinant S contains 2 proline mutations (K986P and V987P), an ablated furin cleavage site, and lacks a transmembrane domain (Figure 1C). A complete picture of the conformational ensemble of S in a native context is essential to understand the dynamic transitions in the SARS-CoV-2 virus lifecycle and accurately depict the epitope landscape in S variants. This is emphasized by the differences between epitope targets from neutralizing antibodies generated by mRNA vaccination and those generated by breakthrough infections.^8^ For example, antibodies targeting the trimer core helix bind both wild-type (WT) and variant recombinant trimers but only neutralize virions displaying WT S.^17^ Furthermore, antibodies targeting the interprotomer NTD-SD1 interface block ACE2 interactions across S variants on virions or cell surfaces but fail to recognize and bind recombinant S. This suggests distinct differences in the overall S conformational landscape and interprotomer contacts in native S from virions.^18,19^

To identify differences between the conformational landscape of recombinant S and S in a native virus environment, we extended HDXMS to probe dynamics of SARS-CoV-2 virus-like particles (VLPs).^20^ VLPs maintain S in a native-like setting, without the risks associated with handling infectious SARS-CoV-2 virus particles. Additionally, the co-expression of auxiliary surface proteins in VLPs allows for the expression of native S without requiring proline substitutions. Our results reveal that while dynamics of recombinant and VLP S are similar for the RBD, large differences in deuterium uptake were observed in regions encompassing interprotomer contacts. Notably, we report a large decrease in deuterium exchange in VLP S at the trimer core interface and the RBD hinge region within SD1 compared to recombinant S. These findings highlight broad differences in the conformational ensemble of native S relative to the recombinant construct. These differences are attributed to engineered modifications and a lack of auxiliary proteins and membrane anchoring in recombinant S.^21^ The observed differences in dynamics between the physiologically relevant VLP S and engineered recombinant S trimers have implications for epitope display in vaccines employing engineered recombinant S. The insights from conformational dynamics of VLP S can inform strategies to mitigate the emergence of breakthrough infections in vaccinated individuals.

## Results

### The conformational landscape of VLP S is distinct from recombinant S with overall decreased exchange in S2

Comparative HDXMS analysis was performed on purified soluble recombinant and VLP S using a linear-IMS-Q-Tof analyzer as described in the methods.^16^ Due to the complex nature of VLP S, we could not obtain sufficient pepsin fragments with high signal-to-noise and therefore used a cyclic-IMS-Q-Tof instrument as described in the methods.^22^ For recombinant S, 277 pepsin fragment peptides were observed, resulting in a coverage of 71.3% (Figure S1A), while for VLP S, 53 pepsin fragment peptides were observed, resulting in a coverage of 44.9% (Figure S1B). To mitigate effects of cold denaturation that promoted the transition from a splayed to tight trimer conformation, both recombinant and VLP S samples were incubated for 3h at 37℃ (Table S1).^13,23^ HDXMS was measured at deuterium exchange times (D_ex_ = 1, 10, and 100 min) for VLP S and a single deuterium exchange timepoint, D_ex_ = 10 min for recombinant D614G S. A longer time course of deuterium exchange for recombinant, WT, and variant S at D_ex_ = 1, 10, 30, 100 min was previously analyzed on the linear ESI-Q-Tof analyzer.^16^ The inherent complexity of VLP S resulted in a distinct pepsin fragmentation profile, favoring longer pepsin fragments on average compared to recombinant S.

Our results demonstrate large decreases in deuterium exchange in peptides encompassing the the S2 subunit (Figure 2A-B and Figure S2). In VLP S, several peptides spanning the S2 (714-730, 799-805, 800-806, 1076-1086, 1163-1171, 1184-1197) showed very low exchange across our experimental timescales (t = 1-100 min) (Figure S3). Additionally, reporter peptide (992-1005) from the core helix that exhibited a progressive decline in exchange across S variants^16^ also showed a substantial decrease in deuterium exchange in VLP S compared to recombinant S (Figure 2C). Interestingly, the deuterium uptake kinetics in S2 peptides from VLP S tend to show a leveling off of deuterium incorporation after 10 min (Figure S4). The observed decreases in exchange in S2 are reflective of stronger interprotomer contacts in the crowded membrane anchored enviroment of VLP S. In contrast, peptides spanning the S1 subdomain exhibited minimal differences in deuterium exchange between VLP and recombinant S at D_ex_ = 10 min (Figure 2 and Figure S2).

**Figure 2.**
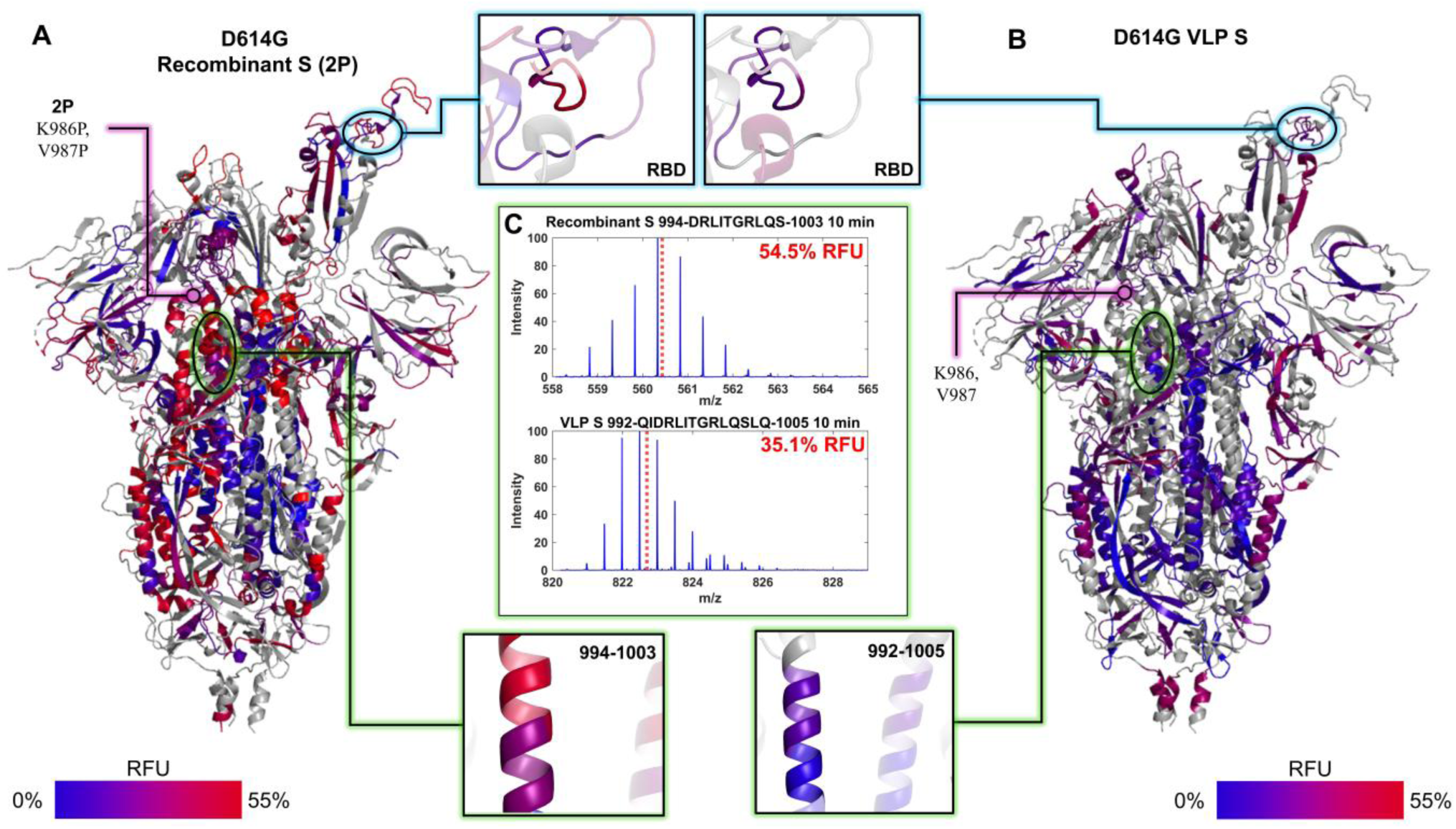
Relative fractional deuterium uptake for D_ex_=10 min mapped onto the S structure (PDB ID: 7T67) for recombinant D614G S **(A)** and for D614G S embedded in VLPs **(B)**. Deuterium exchange is represented by a blue-red gradient up to a maximum 55% uptake. Insets highlight the RBD which shows minimal differences in deuterium uptake (top) and the core helix which shows decreased deuterium uptake in VLP (bottom). 2P substitutions are adjacent to the indicated core helix region indicated in purple. **(C)** Mass spectral envelopes for representative core helical peptide 994-1003 in recombinant S (top) and 992-1005 in VLP S (bottom) after 10 min deuterium exchange. The centroid of each mass spectral envelope is indicated by a red dashed line and the RFU (normalized to 55% maximal uptake) is also denoted in red.

### Altered RBD hinge dynamics in VLP S

Conformational changes between RBD up and RBD down states of S are allosterically coupled to a critical hinge region within the SD1 (Figure 3A-B).^4^ Comparative HDXMS of a peptide (551-570) in this hinge region showed significantly decreased deuterium exchange in VLP S compared to recombinant S (Figure 3C-D). These decreases in exchange are also characterized by the presence of bimodal deuterium exchange mass spectral envelopes in peptides spanning this locus. In the recombinant S, we previously observed the presence of distinct high and low exchanging populations for peptide-spanning residues 553-568.^16^ Spectral deconvolution showed that the low-exchanging population was only observed after 1 min exchange (Figure 3C).^24^ In contrast, a majority (94-97%) of this locus in the VLP S adopts a low exchanging conformation that persists within our experimental labeling timescales (1-100 min) (Figure 3D).

**Figure 3.**
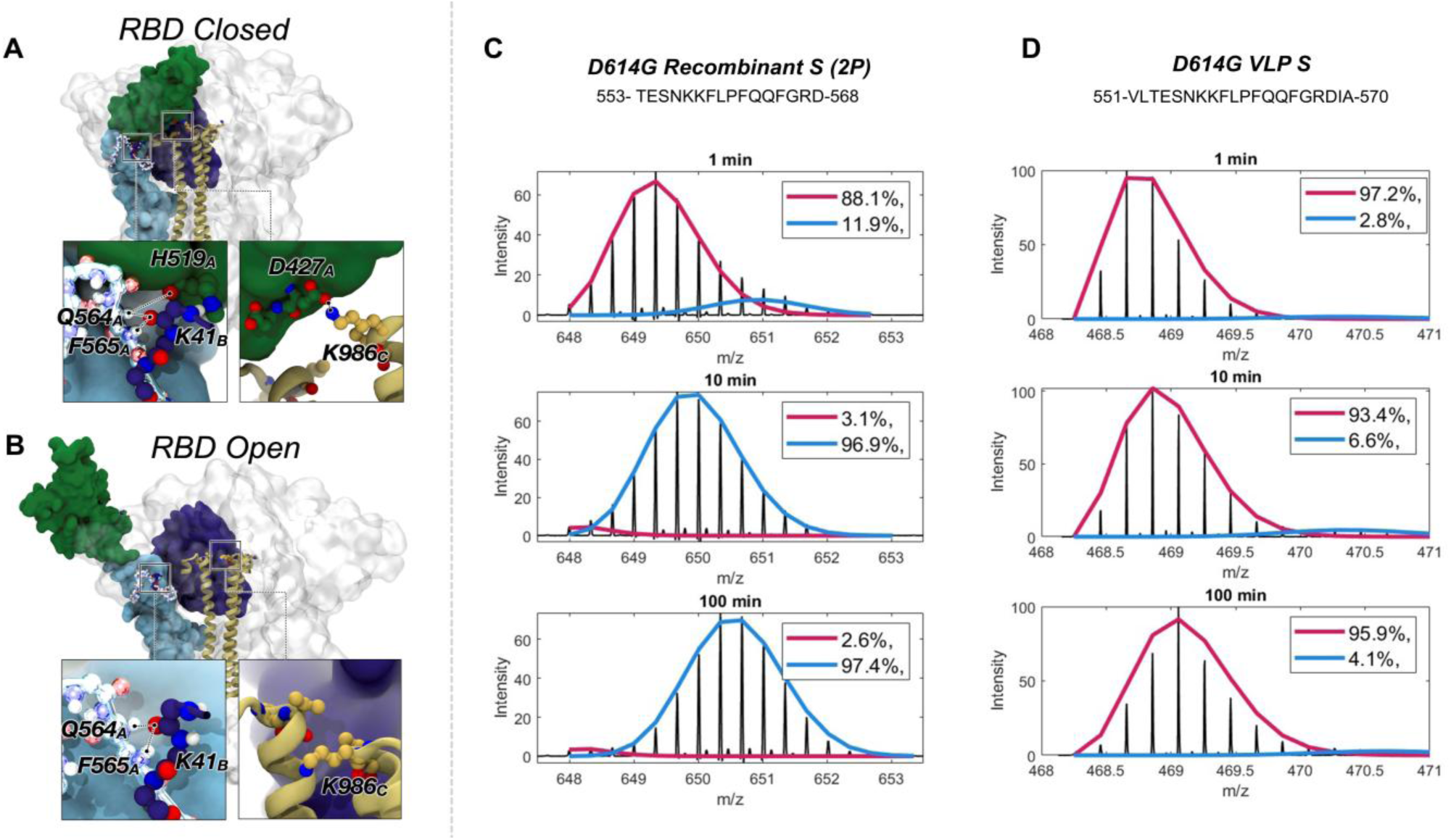
Key inter and intra protomer hydrogen bonds in an RBD closed **(A)** and an RBD open **(B)** state are shown in insets below corresponding S trimer structures. An inter protomer H-bond between K986 and D427 is formed in the RBD closed state. Additional H-bonds are formed between the SD1 hinge of protomer A and NTD of protomer B in both closed and open states. Stacked mass spectral envelopes for undeuterated reference states and after 1, 10, and 100 min exchange (top-bottom) for representative peptide 553-568 in recombinant S **(C)** and representative peptide 551-570 in VLP S **(D)**. Deuterium uptake and mass spectral envelope centroids are indicated in red. Bimodal distributions were deconvolved as a sum of two gaussian distributions using HXexpress3 v39.^23^ The pink curve corresponds to the gaussian fit for the low exchanging population and the blue curve corresponds to the gaussian fit for the high exchanging population.

Key interprotomer contacts at the RBD hinge mediate conformational switching from RBD closed to RBD open. In the closed conformation, the hydrogen bond between D427 in protomer A (D427_A_) and K986 in protomer C (K986_C_) is essential for stabilizing the RBD closed conformation.^25^ Hinge residues Q564_A_ and F565_A_ also interact with H519_A_ and K41_B_ (Figure 3A). When the RBD switches to the open conformation, the salt bridge between D427_A_ and K986_C_ and the hydrogen bond between Q564_A_ and H519_A_ are broken (Figure 3B). 2P substitutions K986P and V987P would disrupt these interprotomer contacts and destabilize the trimer core in recombinant S. The increased propensity for the low exchanging conformation of the hinge reporter peptide in VLP S is reflective of a more intact interprotomer H-bonding network (Figure 3C).^24^ This establishes that membrane-anchored native S in the context of the intact virion is conformationally distinct from engineered recombinant S.

### Trimer cooperativity coordinates RBD opening in VLP S

To assess the interprotomer communication during RBD opening we carried out network analysis on WT and omicron S. This has significant implications for both receptor-mediated allostery and epitope display. Allosteric networks corresponding to RBD opening pathways in both WT and variant S show interprotomer communication. This network bridges all protomers within a trimer and demonstrates the high stability of the S core during RBD opening (Figure 4A-B). In addition to receptor-mediated allostery that is propagated from the RBD to the membrane-spanning regions, our results also highlight the tight cooperativity between individual protomers in VLP S. The crowded membrane environment in VLP enhances the cooperativity of S by stabilizing S2 subunit.

**Figure 4.**
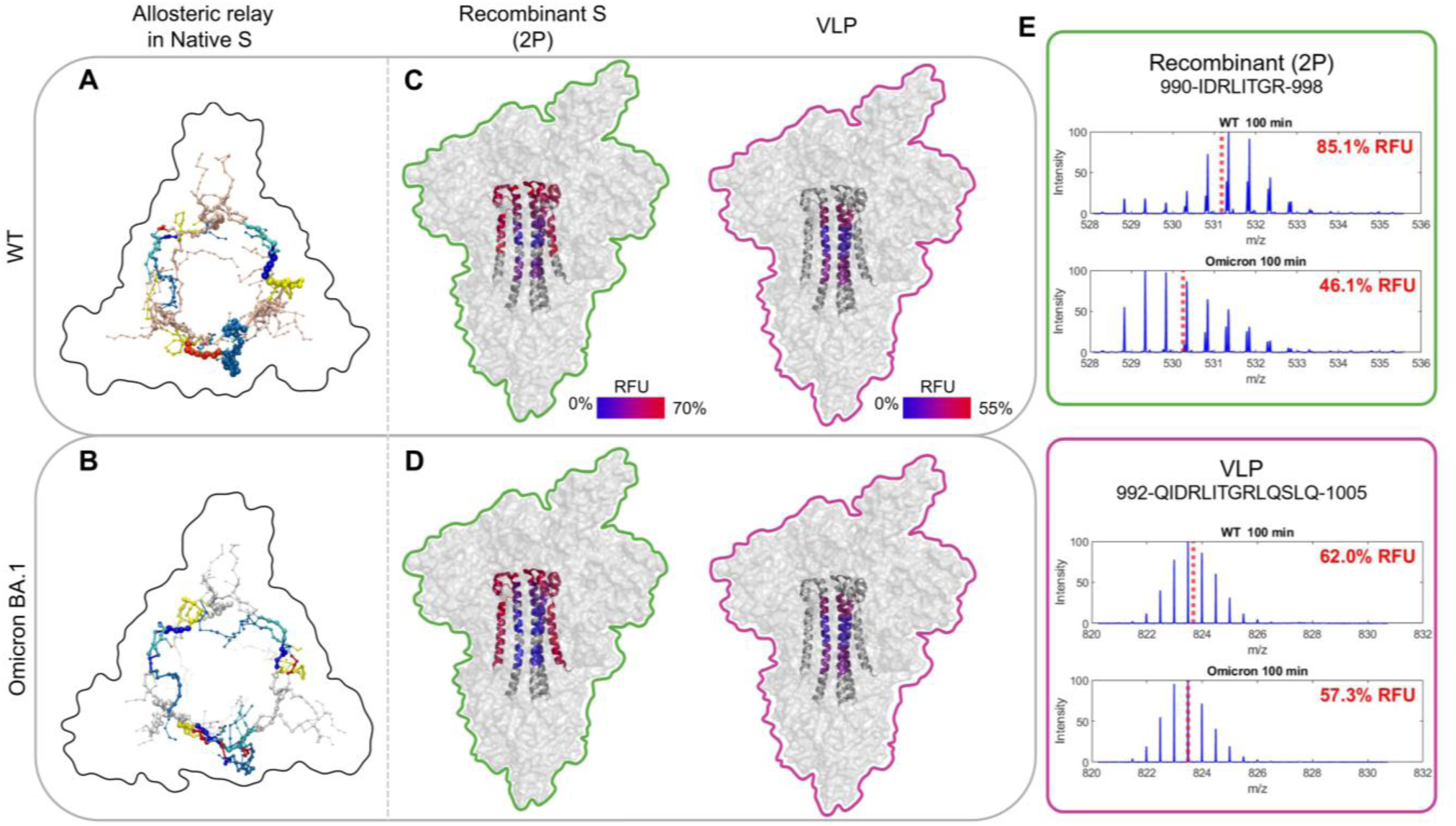
Allosteric relays are mapped onto a top-down view of an outline of the S trimer for WT S **(A)** and omicron S **(B)** showing correlated motions associated with RBD opening. The allosteric relay forms a ring around the trimer connecting residues across protomers. Relative fractional deuterium uptake mapped onto the S core helix (PDB ID: 7T67) for recombinant and VLP (right and left) WT S **(C)** and omicron S **(D)** after 100 min exchange. Deuterium uptake is mapped by a blue-red gradient up to 55% uptake. **(E)** Stacked mass spectral envelopes corresponding to representative CH peptides in recombinant (green) and VLP S (pink) showing protection from deuterium exchange in omicron S (bottom) compared to WT S (top) after 100 min exchange. Relative fractional uptake and centroids for mass spectral envelopes are indicated in red.

### Increased stability of VLP S dampens the magnitude of progressive variant effects on S ensembles

To probe the effects of mutations associated with progressive SARS-CoV-2 variants, we applied comparative HDXMS to VLPs containing WT, D614G, and omicron S. Our results have revealed that the native virion environment enhanced the baseline stability of the S core and plays a key role in RBD transitions (Figure 4C-D). This consequently reduced the impact of variant mutations on magnitude differences in deuterium exchange on VLP S. Emergent variants were previously shown to progressively favor core stability in recombinant S. We observe a similar but lower magnitude stabilization of the core helix in VLP omicron S (Figure 4C-E).

## Discussion

Conformational landscapes of membrane-anchored protein complexes are highly dependent upon quaternary interactions, membrane anchoring, and auxiliary proteins.^26^ This is especially relevant for structural proteins displayed on surfaces of enveloped viruses.^27–30^ Lipidomic profiles of viral membranes have highlighted a correlation between lipid composition and virus infectivity in influenza.^31^ The compositions of lipid envelopes and presence of embedded proteins induce membrane curvature, afford partitioning of specific lipids to create microdomains. These microdomains are likely to modulate conformational dynamics of membrane-anchored viral surface proteins.^32^

Allosteric effects on viral protein conformation are thereby propagated through a combination of quaternary contacts and communication between proteins via the lipid envelope.^33^ In SARS-CoV-2, assembly and budding occur from the ER-Golgi intermediate compartment, where the virion and VLP obtain their lipid envelope.^34^ Lipid-protein interactions of membrane-embedded E, M, and membrane-anchored S, interactions between M and N proteins will, in addition to lipid composition and virion membrane curvature, modulate conformational dynamics of S-embedded in SARS CoV-2 virion membranes to accentuate differences relative to recombinant S.^35,36^ This may be further complemented by S interactions with E, M, N, or adjacent S trimers in the crowded membrane microenvironment of the virion or VLP, which are lacking in S recombinant protein.

HDXMS has shown that the conformation ensemble of S is highly sensitive to perturbations including temperature^13^, receptor binding^14^, and mutations^16^. However, these studies have all been carried out in recombinant S. Recently, HDXMS of eVLPs^37^ has shown local contributions of 2P substitutions, furin ablation, and the presence of a membrane on S conformation.^38^ Here, we have extended these studies to the more physiological native membrane-embedded S as displayed on the virion surface together with all the accessory proteins in the VLP. Our results reveal significant differences in the average conformational ensemble of VLP S. Ensemble differences were primarily observed at interprotomer interfaces stemming from the increased stability of the core trimer interface of the VLP. The increased stability propagated to interprotomer contacts between a hinge in SD1 and the neighboring NTD with significant implications for RBD up/down transitions. Due to the loss of these contacts in recombinant S, RBD up/down transitions in each monomer are likely decoupled from the rest of the trimer. In addition, we demonstrate how membrane anchoring and presence of auxiliary E, M, and N proteins in SARS CoV-2 alter the conformational ensemble baseline of S. Core helix stabilization was observed for both recombinant and VLP S through variant progression.^16^ However, baseline stabilization of VLP S leads to lower magnitude effects.

The observed dynamics and cooperative allostery between protomers make the S trimer a conformationally shifting target for antibody recognition and neutralization. The enhanced expression of the recombinant construct has facilitated the generation of mRNA vaccines that provide significant short-term protection.^39^ However, the impact of the altered conformational landscape observed in recombinant S upon epitope display has not been considered. Despite producing a robust antibody response, mRNA constructs also generate a higher ratio of non-neutralizing antibodies compared to native viral infections.^8^ These differences are especially evident for antibodies that target interprotomer interfaces. For example, antibodies targeting the core helix only effectively bind recombinant S while failing to neutralize variant S virions.^17^ We observe the highest core helix stability in variant S displayed on VLP explaining the lack of core helix epitope accessibility in variant S virions. Additionally, antibodies targeting the hinge in SD1 selectively bind S in native virions.^18,19^ We provide a mechanism for differential antibody binding at this locus since we observe disrupted interprotomer contacts between SD1 and NTD in recombinant S. Together, these results highlight the importance of conformational dynamics of native S for epitope presentation. Overall, the best long-term and cross-variant protection is observed in individuals with both mRNA vaccine-induced and natural immunity.^39,40^ Our work suggests that the natural infection supports the strong short-term immune response of the mRNA vaccine by presenting interprotomer epitopes that are not sampled by the distinct recombinant conformational landscape.

## Methods

### Plasmid constructs

pcDNA3-Membrane, pcDNA3-Nucleoprotein, pCMV-Envelope, and pCMV-Spike plasmids were a gift from Erica Sapphire (The La Jolla Institute of Immunology, La Jolla, CA). Spike variant plasmids pCAGGS-D614G Spike (Addgene #185692, from Marceline Côté) and pCAGGS-BA.1 Spike (Addgene #185452, from Marceline Côté) were obtained from Addgene.

### Expression and purification of SARS-CoV-2 VLPs (WT, D614G, and omicron BA45)

Human embryonic kidney (HEK) 293 cells (American Type Cell Collection, Manassas, VA) were maintained at 37°C and 5% CO_2_ in Dulbecco’s Modified Eagle’s Medium (Thermo Fisher Scientific) supplemented with 10% fetal bovine serum (Biowest USA) and 1% penicillin streptomycin (Thermo Fisher Scientific). HEK293 cells were grown to 70-80% confluency in 100mm round dishes. pcDNA3-M, pcDNA3-N, pCMV-E, and an S protein plasmid (pCMV-wt S, pCAGGS-D614G S, or pCAGGS-BA.1 S) were transfected at a ratio of 1:1:1:2 using a 3:1 ratio of polyethyleneimine hydrochloride (PEI) (Polysciences) to DNA. The DNA and PEI were mixed in Opti-MEM (Fisher Scientific) and incubated at room temperature for 10-15 minutes before being added dropwise to cells. The cells were incubated at 37°C and 5% CO_2_. The cell media was replaced with fresh DMEM supplemented with 10% fetal bovine serum and 1% penicillin streptomycin after five to six hours post-transfection. At 48 hours post-transfection, the media was collected in a falcon tube containing 0.1x HaltTM Protease Inhibitor Cocktail (Thermo Fisher Scientific). The media was clarified by centrifugation at 1000 x *g* for 10 minutes at 4°C, transferred to a new falcon tube, and further clarified by centrifugation at 2000 x *g* for 10 min at 4°C. The supernatant was then loaded onto a 20% sucrose cushion in TNE buffer (50 mM Tris-HCl, 100 mM NaCl, 0.5 mM EDTA, pH = 7.4) and ultracentrifuged at 100,000 x *g* in a Beckman Type 70 Ti rotor for 3 hours at 4°C. VLP pellets were dried, gently resuspended in TNE buffer, and stored at -20°C before shipping. Samples were shipped on ice then flash frozen and stored at -80°C before HDXMS analysis.

### Expression and purification of SARS-CoV-2 recombinant S

Purification of recombinant S was conducted as described in Braet et al. 2023.^16^

### Cryo-EM Screening of VLPs

To visually confirm the presence of S embedded on VLPs, cryo-EM screening was performed. 3.5 uL of sample (D614G VLP) at 0.13 mg/mL was vitrified onto glow discharged copper QuantiFoil 2/1 300 mesh grids. The grid was screened in a 200kV Talos Arctica Microscope at dose rate of 50.88 e/A2 and a pixel size of 0.944 A/px. This revealed highly pleomorphic VLPs ranging from 60-100 nm with multiple S proteins embedded on the VLP surface.

### Deuterium exchange of recombinant S on linear-IMS-Q-Tof

Deuterium labeling of recombinant S was conducted as described in Braet et al. 2023.^16^

### Deuterium exchange of VLP on cyclic-IMS-Q-Tof

Labeling buffer was prepared by diluting 20 X TNE in H_2_O in D_2_O (99.9%). VLP and recombinant samples were incubated at 37 ℃ for 3 hours before labeling to ensure trimer splaying was not observed.^13,23^ 20 μL of sample were added to 40 μL of labeling buffer for a final labeling concentration of 63.3%. Deuterium labeling was conducted for 1, 10, and 100 min at 20°C for VLPs and 10 min at 20°C for recombinant S. After labeling, 20 μL of prechilled quench solution (4 M Guanidinium Hydrochloride, 0.4 M Tris(2-carboxyethyl) phosphine, 2.4 mM n-Dodecyl β-D-Maltoside was added to the 60 μL labeling mixture to bring the reaction to pH 2.5. Reaction conditions are summarized in Table S1.

### LC and Mass spectrometry of recombinant S on linear-IMS-Q-Tof

Deuterated recombinant S samples were analyzed on a Waters Synapt linear-IMS-Q-Tof analyzer as described in Braet et al. 2023.^16^

### LC and Mass spectrometry of VLP on cyclic-IMS-Q-Tof

Deuterated and control samples were digested online at 15 °C using an AffiPro pepsin column (AffiPro, AP-PC-001). All chromatographic components were housed in the UPLC system’s cooling chamber, maintained at 0.0 ± 0.1 °C throughout the measurements. Peptides were trapped and desalted on a VanGuard Pre-Column trap [2.1 mm × 5 mm, ACQUITY UPLC BEH C18, 1.7 μm (Waters, 186002346)] for 3 minutes at a flow rate of 100 μL/min. Elution was achieved using a 5%–35% acetonitrile gradient over 10 minutes at 100 μL/min, with separation on an ACQUITY UPLC HSS T3 column [1.8 μm, 1.0 mm × 50 mm (Waters, 186003535)]. The system operated at an average back pressure of ∼12,950 psi under initial conditions (5% acetonitrile, 95% water, 0.1% formic acid) at 0 °C. Deuterium level determination had an error margin of ± 0.25 Da in this experimental setup. To avoid peptide carryover, a wash solution [1.5 M guanidinium chloride, 0.8% formic acid, and 4% acetonitrile] was injected over the Enzymate column after each analytical run.

Mass spectra were acquired on a Waters SELECT SERIES Cyclic IMS instrument coupled with a UPLC I-Class system and HDX manager. The mass spectrometer was calibrated *via* direct infusion of a major mix solution at 10 μL/min (Waters, 186008113). For post-acquisition mass accuracy correction, glu-fibrinopeptide (Sigma, F3261) was infused at 250 fmol/μL at 5 μL/min as a lock mass. Spectra were recorded over an m/z range of 50–2000 with the following settings: capillary voltage, 3.0 kV; trap collision energy, 4 V; sampling cone, 40 V; transfer CE ramp, 15–50 V; source temperature, 80 °C; and desolvation temperature, 450 °C. Ion mobility separation was performed with a single pass using a sequence of 10 ms injection, 3 ms separation, and 34 ms ejection/acquisition.

### Peptide identification

Peptides of WT and SARS CoV-2 variant S were identified through independent searches of mass spectra from the undeuterated samples. Peptides common to WT and variant S were identified from a database containing the amino acid sequence of WT and D614G S using PROTEIN LYNX GLOBAL SERVER version 3.0 (Waters, Milford, MA) in HDMSE mode for non-specific protease cleavage. Search parameters in PLGS were set to “no fixed or variable modifier reagents” and variable N-linked glycosylation. Deuterium exchange was quantified using DynamX v3.0 (Waters, Milford, MA) with cutoff filters of minimum intensity = 2000, minimum peptide length = 4, maximum peptide length = 25, minimum products per amino acid = 0.2, and precursor ion error tolerance <10 ppm. Three undeuterated replicates were collected for WT and variant S, and the final peptide list includes only peptides that fulfilled the above-described criteria and were identified independently in at least 2 of the 3 undeuterated samples. Deuterium exchange in these peptides was analyzed using DynamX 3.0 with identical parameters described above. To analyze cyclic IMS data, PLGS and DynamX were modified as described in Griffiths et. al. (2024).^41^

### Hydrogen deuterium exchange analysis

The average number of deuterons exchanged in each peptide was calculated by subtracting the centroid mass of the undeuterated reference spectra from each deuterated spectra. Peptides were independently analyzed for quality across technical replicates. Relative deuterium exchange and difference plots were generated by DynamX v3.0. The data for the mass spectra were acquired from DynamX v3.0 and plotted in MATLAB 2024a, The MathWorks Inc, Natick, MA, USA. Relative deuterium exchange plots are reported as RFU which is the ratio of exchanged deuterons to possible exchange deuterons. Reported RFU values for cyclic IMS were normalized to the maximum possible deuterium incorporation without back exchange correction (55% for VLP S, WT, D614G, and omicron and recombinant S, D614G). Reported RFU values for linear IMS were normalized from back exchange corrections described in Braet et al. 2021^16^ (70% for recombinant S WT and omicron). Bimodal deconvolution was carried out by fitting mass spectral envelopes to a sum of two Gaussians using HXexpress3 v39.^24^ The mass spectrometry proteomics data will be deposited to the ProteomeXchange Consortium via the PRIDE partner repository.

### Molecular dynamics simulations and allosteric pathway mapping

In the current work, we have utilized weighted ensemble (WE) molecular dynamics (MD) simulations of spike RBD opening and network analyses performed in conjunction with another related manuscript that is currently submitted and under review. While complete methodological details are provided in that manuscript (submitted, citation to be provided at the time of publication), we will summarize methods here. WE-MD simulations of WT and omicron BA.1 spike proteins were performed similarly to those described by Sztain et al (2021):^25^ We defined a two-dimensional reaction coordinate using (1) the distance between the center of mass of the Cα atoms of the spike RBD central beta-sheets to the center of mass of the Cα atoms of the spike core helices and (2) the root mean square displacement between the RBD β-sheets’ Cα atoms in each simulation frame and the RBD β-sheets’ Cα atoms as seen in PDB ID 6VSB.^6^ All WE-MD simulations were performed with the WESTPA software,^42^ GPU-accelerated Amber MD engine,^43–46^ the CHARMM36m all-atom force field,^47–49^ and the CHARMM-modified TIP3P water model.^50^ Each protein was solvated in a 150 mM NaCl water box of 220 Å x220 Å x220 Å. Hydrogen bonding and salt-bridge analyses from resultant WE-MD simulations were performed via inhouse MDAnalysis scripts.^51,52^ Allosteric networks were calculated from resultant WE-MD RBD opening simulations via the Weighted Implementation of Suboptimal Paths (WISP) method^53^ to calculate the shortest allosteric paths between all residue pairs. NetworkX, using Dijkstra’s algorithm, was used for shortest path computation. Visualization of contacts and allosteric networks was performed using VMD, showing the top 1000 most relevant edges and nodes.^54^

## ASSOCIATED CONTENT

### Supporting Information

HDXMS coverage maps, heatmaps, and experimental design table for D614G recombinant S and WT, D614G, and omicron BA.1 VLP samples. (PDF)

Spreadsheet of bimodal analysis of peptide 553-568 in recombinant D614G S using HXExpress. (.xlsx)

Spreadsheet of bimodal analysis of peptide 551-570 in VLP D614G S using HXExpress. (.xlsx)

## AUTHOR INFORMATION

### Competing interests

The authors declare no competing interests.

### Funding Sources

SARS-CoV-2 VLP research has been supported by the NIH/NIAID (AI169896) to R.V.S. E.J.B. is supported by a NIAID T32 (T32AI148103). Carla Calvó-Tusell acknowledges funding support from Schmidt Sciences, LLC. Simulations were performed on Oracle Cloud, supported by a generous gift from Oracle for Research, and TACC Frontera. We thank Oracle for Research and TACC Frontera for their continued support of our simulations.

## ACKNOWLEDGMENT

We would like to thank Alexandre Gomes and Ignatius Cass from Waters, Milford MA for their assistance in processing cyclic IMS data.

## ABBREVIATIONS

HDXMS: Hydrogen deuterium exchange – mass spectrometry
VLP: virus-like particle
E: SARS-CoV-2 envelope protein
M: SARS-CoV-2 membrane protein
N: SARS-CoV-2 nucleocapsid protein
S: SARS-CoV-2 spike protein
NTD: N-terminal domain
RBD: receptor binding domain
SD1/2: subdomain 1/2
FP: fusion peptide
HR1/2: heptad repeats 1/2
TM: transmembrane domain
CD: cytoplasmic domain
2P: 2 proline mutation at K986P and V987P
HEK: Human embryonic kidney
PEI: polyethyleneimine hydrochloride
PLGS: protein lynx global server
WE: weighted ensemble
MD: molecular dynamics
WISP: weighted implementation of suboptimal paths

## Notes

### Competing Interest Statement

The authors have declared no competing interest.

## REFERENCES

1. Ke, Z. et al. Structures and distributions of SARS-CoV-2 spike proteins on intact virions. Nature 588, 498–502 (2020).

2. Martínez-Flores, D. et al. SARS-CoV-2 Vaccines Based on the Spike Glycoprotein and Implications of New Viral Variants. Front. Immunol. 12, 701501 (2021).

3. Huang, Y., Yang, C., Xu, X.-F., Xu, W. & Liu, S.-W. Structural and functional properties of SARS-CoV-2 spike protein: potential antivirus drug development for COVID-19. Acta Pharmacol. Sin. 41, 1141–1149 (2020).

4. Seow, J. et al. A neutralizing epitope on the SD1 domain of SARS-CoV-2 spike targeted following infection and vaccination. Cell Rep. 40, 111276 (2022).

5. Gobeil, S. M.-C. et al. D614G Mutation Alters SARS-CoV-2 Spike Conformation and Enhances Protease Cleavage at the S1/S2 Junction. Cell Rep. 34, 108630 (2021).

6. Wrapp, D. et al. Cryo-EM structure of the 2019-nCoV spike in the prefusion conformation. Science 367, 1260–1263 (2020).

7. Jackson, C. B., Farzan, M., Chen, B. & Choe, H. Mechanisms of SARS-CoV-2 entry into cells. Nat. Rev. Mol. Cell Biol. 23, 3–20 (2022).

8. Amanat, F. et al. SARS-CoV-2 mRNA vaccination induces functionally diverse antibodies to NTD, RBD, and S2. Cell 184, 3936-3948.e10 (2021).

9. Hsieh, T.-H. S. et al. Resolving the 3D Landscape of Transcription-Linked Mammalian Chromatin Folding. Mol. Cell 78, 539–553.e8 (2020).

10. Pallesen, J. et al. Immunogenicity and structures of a rationally designed prefusion MERS-CoV spike antigen. Proc. Natl. Acad. Sci. U. S. A. 114, E7348–E7357 (2017).

11. Barnes, C. O. et al. SARS-CoV-2 neutralizing antibody structures inform therapeutic strategies. Nature 588, 682–687 (2020).

12. Yang, T.-J., Yu, P.-Y., Chang, Y.-C. & Hsu, S.-T. D. D614G mutation in the SARS-CoV-2 spike protein enhances viral fitness by desensitizing it to temperature-dependent denaturation. J. Biol. Chem. 297, (2021).

13. Costello, S. M. et al. The SARS-CoV-2 spike reversibly samples an open-trimer conformation exposing novel epitopes. Nat. Struct. Mol. Biol. 29, 229–238 (2022).

14. Raghuvamsi, P. V. et al. SARS-CoV-2 S protein:ACE2 interaction reveals novel allosteric targets. eLife 10, e63646 (2021).

15. Yang, Z. et al. SARS-CoV-2 Variants Increase Kinetic Stability of Open Spike Conformations as an Evolutionary Strategy. mBio 13, e03227–21 (2022).

16. Braet, S. M. et al. Timeline of changes in spike conformational dynamics in emergent SARS-CoV-2 variants reveal progressive stabilization of trimer stalk with altered NTD dynamics. eLife 12, e82584 (2023).

17. Silva, R. P. et al. Identification of a conserved S2 epitope present on spike proteins from all highly pathogenic coronaviruses. eLife 12, e83710 (2023).

18. Zhou, D. et al. The SARS-CoV-2 neutralizing antibody response to SD1 and its evasion by BA.2.86. Nat. Commun. 15, 2734 (2024).

19. Liu, L. et al. Antibodies targeting a quaternary site on SARS-CoV-2 spike glycoprotein prevent viral receptor engagement by conformational locking. Immunity 56, 2442–2455.e8 (2023).

20. Plescia, C. B. et al. SARS-CoV-2 viral budding and entry can be modeled using BSL-2 level viruslike particles. J. Biol. Chem. 296, 100103 (2021).

21. Amanat, F. et al. Introduction of Two Prolines and Removal of the Polybasic Cleavage Site Lead to Higher Efficacy of a Recombinant Spike-Based SARS-CoV-2 Vaccine in the Mouse Model. mBio 12, e02648–20 (2021).

22. Giles, K. et al. A Cyclic Ion Mobility-Mass Spectrometry System. Anal. Chem. 91, 8564–8573 (2019).

23. Edwards, R. J. et al. Cold sensitivity of the SARS-CoV-2 spike ectodomain. Nat. Struct. Mol. Biol. 28, 128–131 (2021).

24. Tuttle, L. M. et al. Rigorous Analysis of Multimodal HDX-MS Spectra. J. Am. Soc. Mass Spectrom. 36, 416–423 (2025).

25. Sztain, T. et al. A glycan gate controls opening of the SARS-CoV-2 spike protein. Nat. Chem. 13, 963–968 (2021).

26. Groves, J. T. & Kuriyan, J. Molecular mechanisms in signal transduction at the membrane. Nat. Struct. Mol. Biol. 17, 659–665 (2010).

27. Lim, X.-X. et al. Epitope and Paratope Mapping Reveals Temperature-Dependent Alterations in the Dengue-Antibody Interface. Structure 25, 1391–1402.e3 (2017).

28. Lim, X.-X. et al. Human antibody C10 neutralizes by diminishing Zika but enhancing dengue virus dynamics. Cell 184, 6067–6080.e13 (2021).

29. Kant, R. et al. Small Molecule Assembly Agonist Alters the Dynamics of Hepatitis B Virus Core Protein Dimer and Capsid. J. Am. Chem. Soc. 146, 28856–28865 (2024).

30. Casalino, L. et al. Breathing and Tilting: Mesoscale Simulations Illuminate Influenza Glycoprotein Vulnerabilities. ACS Cent. Sci. 8, 1646–1663 (2022).

31. Ivanova, P. T. et al. Lipid composition of viral envelope of three strains of influenza virus – not all viruses are created equal. ACS Infect. Dis. 1, 399–452 (2015).

32. Teissier, É. & Pécheur, E.-I. Lipids as modulators of membrane fusion mediated by viral fusion proteins. Eur. Biophys. J. 36, 887–899 (2007).

33. Venkatakrishnan, V., Braet, S. M. & Anand, G. S. Dynamics, allostery, and stabilities of whole virus particles by amide hydrogen/deuterium exchange mass spectrometry (HDXMS). Curr. Opin. Struct. Biol. 86, 102787 (2024).

34. Scherer, K. M. et al. SARS-CoV-2 nucleocapsid protein adheres to replication organelles before viral assembly at the Golgi/ERGIC and lysosome-mediated egress. Sci. Adv. 8, eabl4895 (2022).

35. Dutta, M., Su, Y., Plescia, C. B., Voth, G. A. & Stahelin, R. V. The SARS-CoV-2 nucleoprotein associates with anionic lipid membranes. J. Biol. Chem. 300, 107456 (2024).

36. Dutta, M. et al. Direct lipid interactions control SARS-CoV-2 M protein conformational dynamics and virus assembly. BioRxiv Prepr. Serv. Biol. 2024.11.04.620124 (2024) doi:10.1101/2024.11.04.620124.

37. Hoffmann, M. A. G. et al. ESCRT recruitment to SARS-CoV-2 spike induces virus-like particles that improve mRNA vaccines. Cell 186, 2380–2391.e9 (2023).

38. Shoemaker, S. R. et al. The Interplay of Furin Cleavage and D614G in Modulating SARS-CoV-2 Spike Protein Dynamics. 2025.01.27.635166 Preprint at 10.1101/2025.01.27.635166 (2025).

39. SARS-CoV-2 humoral and cellular immunity following different combinations of vaccination and breakthrough infection - PubMed. https://pubmed.ncbi.nlm.nih.gov/36732523/.

40. Hopkins, S., Hall, V. & Klenerman, P. Protection against SARS-CoV-2 after Vaccination and Previous Infection. Reply. N. Engl. J. Med. 386, 2535 (2022).

41. Griffiths, D. et al. Cyclic Ion Mobility for Hydrogen/Deuterium Exchange-Mass Spectrometry Applications. Anal. Chem. 96, 5869–5877 (2024).

42. Zwier, M. C. et al. WESTPA: an interoperable, highly scalable software package for weighted ensemble simulation and analysis. J. Chem. Theory Comput. 11, 800–809 (2015).

43. Case, D. A. et al. AmberTools. J. Chem. Inf. Model. 63, 6183–6191 (2023).

44. D.A. Case, H.M. Aktulga, K. Belfon, I.Y. Ben-Shalom, J.T. Berryman, S.R. Brozell, D.S. Cerutti, T.E. Cheatham, III, G.A. Cisneros, V.W.D. Cruzeiro, T.A. Darden, N. Forouzesh, M. Ghazimirsaeed, G. Giambaşu, T. Giese, M.K. Gilson, H. Gohlke, A.W. Goetz, J. Harris, Z. Huang, S. Izadi, S.A. Izmailov, K. Kasavajhala, M.C. Kaymak, A. Kovalenko, T. Kurtzman, T.S. Lee, P. Li, Z. Li, C. Lin, J. Liu, T. Luchko, R. Luo, M. Machado, M. Manathunga, K.M. Merz, Y. Miao, O. Mikhailovskii, G. Monard, H. Nguyen, K.A. O’Hearn, A. Onufriev, F. Pan, S. Pantano, A. Rahnamoun, D.R. Roe, A. Roitberg, C. Sagui, S. Schott-Verdugo, A. Shajan, J. Shen, C.L. Simmerling, N.R. Skrynnikov, J. Smith, J. Swails, R.C. Walker, J. Wang, J. Wang, X. Wu, Y. Wu, Y. Xiong, Y. Xue, D.M. York, C. Zhao, Q. Zhu, and P.A. Kollman. Amber 2024. University of California (2024).

45. Salomon-Ferrer, R., Götz, A. W., Poole, D., Le Grand, S. & Walker, R. C. Routine Microsecond Molecular Dynamics Simulations with AMBER on GPUs. 2. Explicit Solvent Particle Mesh Ewald. J. Chem. Theory Comput. 9, 3878–3888 (2013).

46. Le Grand, S., Götz, A. W. & Walker, R. C. SPFP: Speed without compromise—A mixed precision model for GPU accelerated molecular dynamics simulations. Comput. Phys. Commun. 184, 374–380 (2013).

47. Huang, J. & MacKerell Jr, A. D. CHARMM36 all-atom additive protein force field: Validation based on comparison to NMR data. J. Comput. Chem. 34, 2135–2145 (2013).

48. Huang, J. et al. CHARMM36m: an improved force field for folded and intrinsically disordered proteins. Nat. Methods 14, 71–73 (2017).

49. Guvench, O., Hatcher, E., Venable, R. M., Pastor, R. W. & MacKerell, A. D. Jr. CHARMM Additive All-Atom Force Field for Glycosidic Linkages between Hexopyranoses. J. Chem. Theory Comput. 5, 2353–2370 (2009).

50. Mark, P. & Nilsson, L. Structure and Dynamics of the TIP3P, SPC, and SPC/E Water Models at 298 K. J. Phys. Chem. A 105, 9954–9960 (2001).

51. Michaud-Agrawal, N., Denning, E. J., Woolf, T. B. & Beckstein, O. MDAnalysis: A toolkit for the analysis of molecular dynamics simulations. J. Comput. Chem. 32, 2319–2327 (2011).

52. Gowers, R. J. et al. MDAnalysis: A Python Package for the Rapid Analysis of Molecular Dynamics Simulations. scipy (2016) doi:10.25080/Majora-629e541a-00e.

53. Van Wart, A. T., Durrant, J., Votapka, L. & Amaro, R. E. Weighted Implementation of Suboptimal Paths (WISP): An Optimized Algorithm and Tool for Dynamical Network Analysis. J. Chem. Theory Comput. 10, 511–517 (2014).

54. Humphrey, W., Dalke, A. & Schulten, K. VMD: visual molecular dynamics. J. Mol. Graph. 14, 33–38, 27–28 (1996).

